# Targeted Proteomics Reveals Quantitative Differences in Low Abundance Glycosyltransferases of Patients with Congenital Disorders of Glycosylation

**DOI:** 10.1101/2020.09.15.291732

**Authors:** Roman Sakson, Lars Beedgen, Patrick Bernhard, Keziban M. Alp, Nicole Lübbehusen, Ralph Röth, Beate Niesler, Matthias P. Mayer, Christian Thiel, Thomas Ruppert

**Author notes:** equal contribution (Roman Sakson, Lars Beedgen).

## Abstract

Protein glycosylation is essential in all domains of life and its mutational impairment in humans can result in severe diseases named Congenital Disorders of Glycosylation (CDGs). Studies on molecular level are however challenging, because many glycosyltransferases in the endoplasmic reticulum (ER) are low abundance membrane proteins. We established a comprehensive multiple reaction monitoring (MRM) assay to quantify most human glycosyltransferases involved in the processes of *N*-glycosylation,*O*- and *C*-mannosylation in the ER. To increase reproducibility, a membrane protein fraction of isotopically labeled HEK 293T cells was used as an internal standard. With this internal standard the MRM assay is easily transferable between laboratories. 22 glycosyltransferases could be reliably quantified from whole cell lysates of HEK 293T cells, HeLa cells and skin fibroblast cell lines. We then analyzed fibroblasts derived from CDG type I patients with mutations in the *ALG1*,*ALG2* or *ALG11* gene. Mutations in *ALG1* or *ALG2* gene strongly reduced the levels of the ALG1 and ALG2 protein, respectively. In contrast, the levels of all other glycosyltransferases remained unchanged, which was unexpected given evidence that the ALG1, ALG2 and ALG11 proteins form a stable complex. This study describes an efficient workflow for the development of MRM assays for low abundance proteins, establishes a ready-to-use tool for the comprehensive quantification of ER-localized glycosyltransferases and provides new insight into the organization of disease-relevant glycosylation processes.

---

Protein glycosylation is an essential modification that is conserved in all kingdoms of life. *N-*glycosylation is the most common form of glycosylation in mammals. At least 13 different asparagine-linked glycosylation (ALG) glycosyltransferases and the putative flippase RFT1 are required for the production of the dolichol-linked N-glycan precursor in the endoplasmic reticulum (ER), before the precursor is subsequently transferred to the nascent chain by the oligosaccharyltransferase (OST) complex^1^ (Figure 1a). Protein *O-*mannosylation and protein *C-*mannosylation also start in the ER and are mediated by the protein *O-*mannosyltransferases (POMT1 and POMT2, TMTC1, TMTC2, TMTC3 and TMTC4) and the *C-*mannosyltransferases (DPY19L1, DPY19L3 and DPY19L4).^2, 3^ All three forms of glycosylation require nucleotide- or dolichol phosphate-activated mannose, the production of which requires, among others, the PMM2, DPM1 and DPM3 enzymes.^4^ It is expected that such complex pathways are organized in protein complexes. There is, however, only indirect evidence for higher-ordered structures in humans^5^ due to a lack of tools to precisely quantify most of these low abundance membrane proteins from cellular material. Only for ALG2 and ALG11 co-immunoprecipitation studies in yeast suggested that they form heteromeric complexes with ALG1, which itself can form oligomers.^6^ Because protein glycosylation is an essential modification, severe developmental defects named Congenital Disorders of Glycosylation (CDGs) are linked to mutations that affect the different protein glycosylation processes in humans.^7^ Most of the CDG subtypes known to date affect N-glycosylation processes.^8^

**Figure 1.**
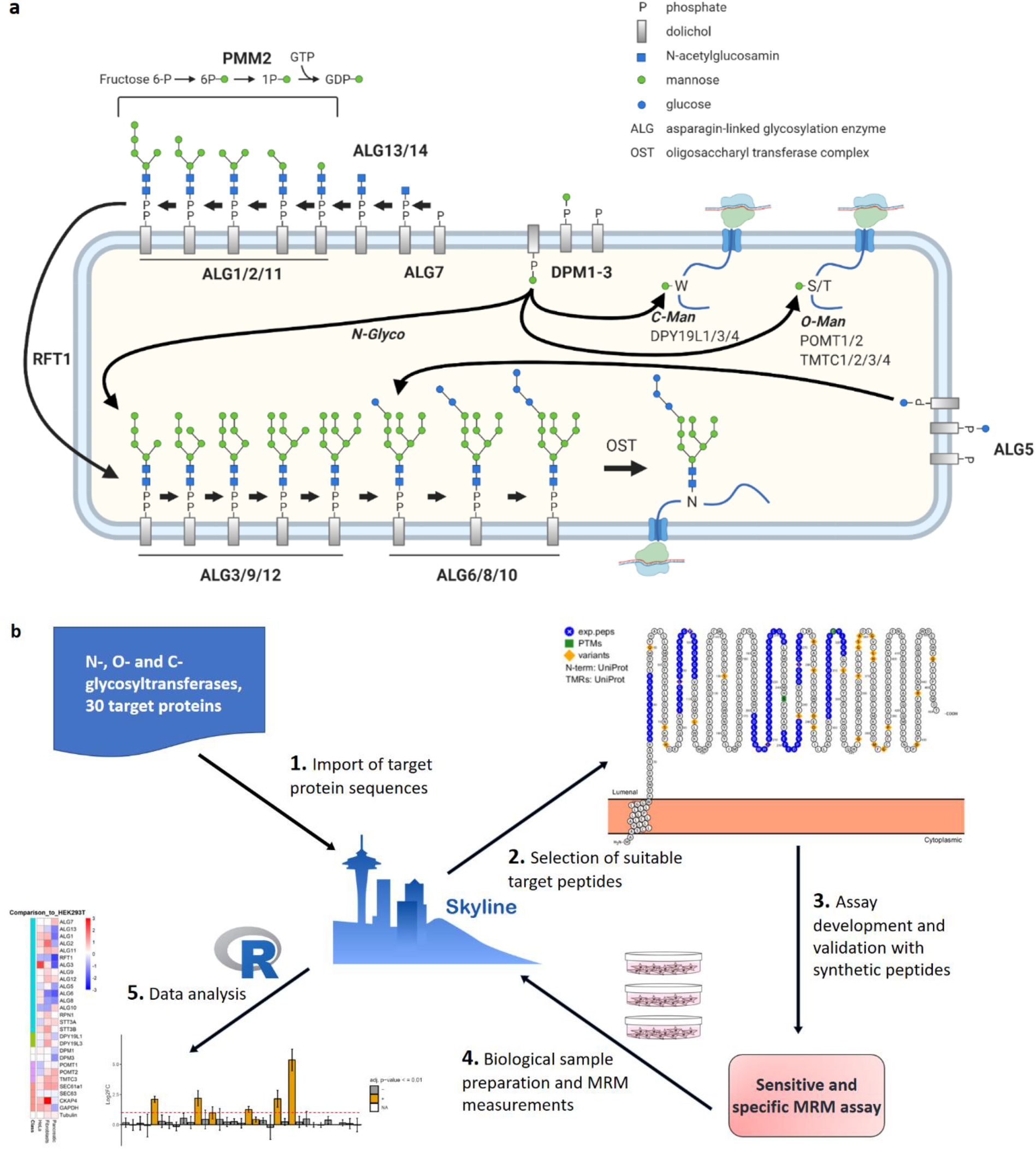
Overview of ER-resident glycosyltransferases and the overall workflow employed in this study. (a) Protein glycosylation in the ER. *N-*glycosylation starts with the production of the *N-*glycan precursor by the addition of *N-*acetyl glucosamine, glucose and mannose residues to dolichol phosphate. Functional units, such as ALG1, ALG2 and ALG11, are written as ALG1/2/11. Dol-P-Man is a substrate for *N-*glycosylation, but also for *O-*mannosylation (*O-*Man) and for *C-*mannosylation (*C-*Man). (b) Development of the MRM method is based on the Skyline software, starting with peptide selection and filtering using the implemented Protter software. Optimization of different parameters is also done with Skyline. As a result, all information necessary to use the method is included in a single Skyline file, which is also used to enable data acquisition of the biological samples. Finally, data interpretation is facilitated by R-scripts that use exported csv-files from the Skyline document as input.

Targeted proteomics strategies like multiple reaction monitoring (MRM, also referred to as selected reaction monitoring or SRM) and parallel reaction monitoring have emerged as the method of choice for the high-confidence identification and quantification of proteins in complex biological samples.^9, 10^ Its routine application is, however, hampered by the complicated and time-consuming setup of such a method. On the other hand, once established in a robust way, such methods can be easily transferred between laboratories and used by the scientific community.^11, 12^ Here, we developed a set of MRM assays for 26 glycosyltransferases involved in ER-based glycosylation processes. In addition, we provide MRM assays for PMM2, RFT1, the auxiliary OST subunit RPN1 and the translocation protein SEC63 as a normalization control. 22 of these enzymes could be precisely quantified from whole cell lysates of skin fibroblasts. We could show that the almost complete loss of ALG1 and ALG2 in primary fibroblasts from CDG patients did not lead to changes in transcript or protein abundance of any of the other ALG enzymes, which means that there is no fail-safe mechanism for the production of the *N-*glycan precursor.

## Experimental Section

### Cell Culture

The study was performed in accordance with the Declaration of Helsinki and approved by the Ethics Committee of the Medical Faculty Heidelberg (S-904/2019). Written informed consent was obtained from the patients’ parents for laboratory analysis of patient material. The cells were maintained at 37 °C in a humidified atmosphere under 5 % CO_2_. Patient and control fibroblasts were cultured in Dulbecco’s modified Eagle’s medium (high glucose; Life Technologies) supplemented with 10 % FCS (PAN Biotech), 100 U/mL penicillin and 100 μg/mL streptomycin. The medium was replaced every 72 h. For extract preparation, 80 – 90 % confluent cell monolayers of a T75 flask were washed with ice-cold PBS and harvested using a cell scraper. Unlabeled HEK 293T and HeLa cells were cultured in Dulbecco’s modified Eagle’s medium (Thermo Fisher Scientific) supplemented with 10 % FBS (Thermo Fisher Scientific), 100 U/mL penicillin and 100 μg/mL streptomycin. For the preparation of cell lysates, cell monolayers (80-90 % confluency) of a T75 flask were washed with PBS and harvested using 0.25 % (v/v) trypsin-EDTA solution. All cell pellets were shock-frozen in liquid nitrogen and stored at −80 °C until cell lysis. For further information about used cell lines, see Table S1.

For metabolic labeling, HEK 293T cells were cultured in DMEM for SILAC (Thermo Fisher Scientific) supplemented with 10 % (v/v) FBS, 100 U/mL penicillin and 100 μg/mL streptomycin, 1 % (v/v) GlutaMAX^®^ (Thermo Fisher Scientific), 0.55 mM [^13^C_6_, ^15^N_4_]-arginine and 0.82 mM [^13^C_6_, 15N2]-lysine (Cambridge Isotopes) for at least seven passages to promote incorporation of isotope-labeled amino acids.^13^ Cell monolayers (80-90 % confluency) of 150 mm cell culture dishes were harvested and stored as unlabeled HEK 293T cells. The preparation of the membrane protein fraction was performed as described (see supporting Materials and Methods for details).^14^ The resulting pellet was resuspended in RIPA buffer, aliquoted, snap-frozen in liquid nitrogen and stored at −80 °C.

### RNA preparation and Nanostring / nCounter^®^ analysis

Total RNA was extracted from approx. 8×105 cells with the RNeasy mini kit (Qiagen) in combination with the QIAshredder system (Qiagen) according to the manufacturer’s protocol. RNA was eluted in 30 μl nuclease-free H2O. RNA quantity and quality were assessed using NanoDrop 2000 (Thermo Fisher Scientific), the Agilent Bioanalyzer 2100 (TotalRNA Nanokit) and the Qubit™ Fluorometer (Qubit RNA HS Kit; Invitrogen). 50 ng of total RNA were used per hybridization reaction. Transcript abundance of 18 glycosyltransferase genes and control genes was determined using an nCounter^®^ SPRINT Profiler (NanoString Technologies) at the nCounter Core Facility of the University of Heidelberg. Probe design for the glycosyltransferase genes and reference genes are given on request. For each patient and control fibroblast cell line, 3 - 4 different cell pellets were analyzed (for raw data, see Table S2). Data were analyzed according to the manufacturer’s guidelines using nSolver Analysis Software 4.0, including calculation of false discovery rate. Housekeeping genes *C1ORF43* and *SNRPD3* were used for normalization. All results were plotted with Prism 8.3.0.

### MRM Assay Development and Refinement

For target proteins, all unique tryptic peptides without missed cleavages and with no methionine in the sequence (unless no other option available) between 7 and 21 amino acids in length were considered. Whenever more than 10 peptide candidates were available based on protein sequence, we used our own DDA data and public repositories^15, 16^ as well as the predictor tool CONSeQuence^17^ to make the final selection (for more details see supporting Materials and Methods). Heavy [^13^C_6_ ^15^N_2_]-lysine-labeled and [^13^C_6_ ^15^N_4_]-arginine-labeled peptides were ordered from JPT and SynPeptide and contained carbamidomethyl modified cysteine residues, if applicable. Peptides were solubilized accordingly to manufacturer’s instructions and analyzed using a Waters nanoACQUITY UPLC System equipped with a trapping column (Waters, Symmetry C18; 2 cm length, 180 μm inner diameter, 5 μM C18 particle size, 100 Å pore size) and an analytical column (Waters, M-Class Peptide BEH C18; 25 cm length, 75 μm inner diameter, 1.7 μM C18 particle size, 130 Å pore size). 1 pmol per peptide was trapped for 7 min at a flow rate of 10 μL/min with 99.4 % of buffer A (1 % v/v ACN, 0.1 % v/v FA) and 0.6 % of buffer B (89.9 % v/v ACN, 0.1 % v/v FA) and separated at the analytical column temperature of 60 °C, with a flow rate of 300 nL/min and a 37 min linear gradient of buffer B in buffer A from 3 to 37 % B, followed by washing and reconditioning of the column to 3 % B. The nanoACQUITY UPLC System was coupled online to an ESI-QTrap 5500 via a NanoSpray III Source (both Sciex). Uncoated pre-cut emitters (Silica Tip, 20 μm inner diameter, 10 μm tip inner diameter, New Objectives, Woburn, MA) and a voltage of approximately 2.6 kV were applied for electrospray ionization. MRM signals were validated using the MRM-initiated detection and sequencing (MIDAS) approach as described^18^ with minor adaptations (see supporting Materials and Methods for details). Data analysis was performed in Skyline.^19^ Peptides with very weak or ambiguous chromatograms were excluded from further analysis. To enable scheduled MRM analysis and easy assay transfer between LC-MS systems and laboratories, indexed retention times (iRTs) were calculated for all peptides using a 10 out of 11 set of widely used peptide iRT calibrators^20^ (all but LGGNEQVTR). Based on MS/MS spectra from the MIDAS measurements, 3-4 most intense transitions per precursor ion were selected (preferably y-ion series, starting from at least y3 or higher). Collision energies were optimized for all transitions in Skyline as published before.^21^ All synthetic peptides were spiked into a complex mixture of three different human whole cell lysates to verify detectability in biological matrices.

### Sample preparation for LC*-*MRM

Sample preparation and LC*-*MS measurements were performed at two different sites (Heidelberg and Freiburg), differences in the protocol are indicated. Fibroblast cells were lysed in 100 μL RIPA buffer (Thermo Fisher Scientific) supplemented with 1 % protease inhibitor (Roche) mix and 1 μL benzonase (250 units/μL, Merck) for 30 min on ice. The cell lysate was passed 10-times through a 20G needle and then centrifuged at 13,000 rpm at 4 °C for 30 min. For HEK 293T and HeLa cells, the total amount of benzonase used was 750 units and needle treatment was omitted. Protein concentration was determined by DC protein Bio*-*Rad kit. Cell lysates were shock-frozen in liquid nitrogen and stored at −80 °C till further analysis. Sample amounts corresponded to approx. 31 – 40 μg of total protein, 20 – 28 % (w/w) of which consisted of the internal standard, depending on the particular experiment. Later, all MRM signals were normalized to the internal standards and, in addition, to SEC63 to account for possible pipetting and protein concentration determination errors. After spike-in, proteins were precipitated using a mixture of methanol and chloroform according to a published protocol.^22^

For in*-*solution digestion, pellets of precipitated proteins were resuspended in 20 μL of urea buffer (8 M urea, Carl Roth, 100 mM NaCl, Applichem, in 50 mM triethylammonium bicarbonate (TEAB), pH 8.5, Sigma-Aldrich). Cysteine thiols were subsequently reduced and alkylated by adding Tris(2-carboxyethyl)phospin (Carl Roth) to a final concentration of 10 mM and 2-Chloroacetamide (Sigma-Aldrich) to a final concentration of 40 mM. The solution was incubated for 30 min at RT. Sample pre-digestion was performed adding 2.5 μg Lysyl Endopeptidase^®^ (Wako Chemicals) and incubating for 4 hours at 37 °C. After diluting the urea concentration to 2 M by adding 50 mM TEAB buffer, 1 μg of trypsin (Thermo Fisher Scientific) was added and samples were incubated for 16-17 hours at 37 °C. Digestion process was stopped by reducing the pH < 2 through the addition of trifluoroacetic acid (TFA, Sigma-Aldrich) to a final concentration of 0.4 – 0.8 % (v/v) and centrifuged for 10 min at 2500 g at RT. Each sample was divided in half to minimize material loss before desalting. In Heidelberg, samples were desalted with C18 StageTips containing 3 disks of Empore C18 material (3M).^23^ In Freiburg, samples were desalted with Hypersep C18 SpinTips (Thermo Fisher Scientific) according to the manufacturer’s instructions. Samples were dried in a vacuum centrifuge and stored at −20 °C until LC*-*MS analysis.

### LC*-*MRM Analysis

Peptides were resolubilized in 20 % ACN / 0.1 % (v/v) TFA and incubated for 5 min at RT, then diluted 10-fold with 0.1 % of TFA to gain a final concentration of 2 % ACN. Stock solutions were stored at −20 °C. Approx. 8 to 9 μg of tryptic peptides were analyzed per LC*-*MRM injection. In Heidelberg, LC*-*MRM analysis was performed using the same LC*-*MS setup and gradient as described above for the assay development. In Freiburg, LC*-*MRM analysis was performed using a nanoflow LC system, Easy-nLC II (Proxeon Biosystems, now Thermo Scientific) equipped with a trapping column (Fused Silica Capillary; 3 cm length, 100 μm inner diameter, VICI Jour, Schenkon, Switzerland) and an analytical column (Self-Pack PicoFrit Column; 40 cm length, 75 μm inner diameter, New Objective, Woburn, MA) both in*-*house packed^24^ with C18 particles (Dr. Maisch, ReproSil-Pur 120 C18-AQ; 3 μm C18 particle size, 120 Å pore size). Samples were trapped at 220 bar with 100 % buffer A (0.1 % v/v FA) and separated using a reversed phase C18 column with the analytical column temperature set at 60 °C and at a flow rate of 250 nL/min. A multistep binary gradient was used for separation: linear gradient of buffer B (50 % v/v ACN, 0.1 % v/v FA) in buffer A from 8 to 56 % B in 60 min, followed by washing and reconditioning of the column to 5 % B. The Easy-nLC II system was coupled online to an ESI-TSQ Vantage triple quadrupole mass spectrometer via a Nanospray Flex Ion source (both Thermo Scientific). The analytical column contains an integrated uncoated pre-cut emitter (Silica Tip, 10 μm tip inner diameter, New Objective, Woburn, MA) and a voltage of approximately 2.5 kV were applied for electrospray ionization. All samples were measured in a randomized order. For a detailed overview of samples and biological replicates used for individual experiments, see Table S3.

### LC-MRM Data and Statistical Analysis

All MRM data were processed and analyzed using Skyline 64-bit version 20.1.0.155 and two in-house written R-scripts, which are available on GitHub including input and output files (https://github.com/alpmerve/MRMStaR, accessed September 15, 2020). Data sets were imported into Skyline and signals were briefly reviewed manually, using the Peak Areas and Retention Time visualization features. We ensured via the Results Grid that all reference dot-product (rdotp) correlations for ratios of the observed transition intensities between corresponding light and heavy precursor pairs were above 0.9. All transitions, for which Skyline reported “Coeluting” as false or “Fwhm Degenerate” as true were manually inspected. Retention times corresponding to the beginning and the end of peak integration boundaries as well as to its apex were used to calculate the symmetry score for each transition. Precursors with very high as well as very low or N/A values were manually inspected. Then, all data was exported from Skyline using the provided Document Grid report template and imported in csv-format into R version 3.6.1 (2019-07-05) “Action of the Toes”.^25^ Briefly, all quantitative and non-truncated transitions were summed up across proteins and log2 light/heavy ratios were computed, to account for technical variance. In addition, all samples were normalized to its respective Sec63 protein level as a general ER marker membrane protein to account for biological variance. Then, pairwise group comparisons were performed using linear regression models between protein ratios from different conditions and controls. Computed p-values were adjusted using the Benjamini and Hochberg multiple test correction.^26^ R code used for pairwise group comparisons was inspired by the R package MSstats.^27^ All results were plotted with Prism 8.3.0.

## Results

### MRM assay for the Quantitative Analysis of ER-based Glycosyltransferases

The major goal of this study was to develop a standardized, quantitative method for the members of the glycosylation machinery in the human ER (Figure 1b for workflow overview). The complete list of 30 target proteins is given in Table 1. Because most of our targets were low abundance transmembrane proteins, we started with up to 13 candidate peptides (median = 9) per protein for method development using the freely available Skyline software suite for targeted proteomics. Peptides were examined for previously reported natural variants and modifications using Protter^28^ via a Skyline plugin. To ensure specificity and achieve the highest sensitivity of the assay, instrument parameters were optimized using commercially available heavy labeled synthetic peptides for all selected peptide candidates. We determined indexed retention times (iRTs)^20^ for all peptides present in the MRM assay, which greatly facilitated the transfer of the method to a different laboratory. Out of 265 ordered synthetic peptides, 211 passed our quality control criteria and were tested in a representative background matrix consisting of a mixture of three different human cell lysates. This procedure led to a ready-to-use MRM assay consisting of 181 peptides for 30 proteins (Tables S4 and S5) with all required information and settings stored within one single Skyline document.

**Table 1.** Overview of target proteins, for which MRM assays were developed in this study^a^.

| Protein name | Uniprot accession | Protein name | Uniprot accession |
| --- | --- | --- | --- |
| PMM2 | O15305 | ALG8 | Q9BVK2 |
| DPM1 | O60762 | ALG10 | *Q5BKT4* |
| DPM3 | Q9P2X0-2 | RPN1 | P04843 |
| ALG7 | Q9H3H5 | STT3A | Q8TCJ2 |
| ALG13 | Q9NP73-2 | STT3B | P46977 |
| ALG14 | Q96F25 | POMT1 | Q9Y6A1-2 |
| ALG1 | Q9BT22 | POMT2 | Q9UKY4 |
| ALG2 | Q9H553 | TMTC1 | Q8IUR5 |
| ALG11 | Q2TAA5 | TMTC2 | Q8N394 |
| RFT1 | Q96AA3 | TMTC3 | Q6ZXV5 |
| ALG3 | Q92685 | TMTC4 | Q5T4D3 |
| ALG9 | Q9H6U8 | DPY19L1 | *NP_001353602.1* |
| ALG12 | Q9BV10 | DPY19L3 | Q6ZPD9 |
| ALG5 | Q9Y673 | DPY19L4 | Q7Z388 |
| ALG6 | Q9Y672 | SEC63 | Q9UGP8 |

<sup>a</sup>ALG10 peptides were shared with ALG10B (Q5I7T1); for DPY19L1 the given NCBI identifier was used instead of the UniProtKB entry Q2PZI1

To enable automated data processing and analysis, we wrote two R-scripts that operate with the output from the Skyline software and that are openly available online.

A membrane fraction of heavy labeled HEK 293T cells was used as an internal SILAC standard for all MRM measurements with biological samples. Most of the target proteins were 8-fold enriched (Figure S1). Successful enrichment also had the beneficial effect of reducing the complexity of the internal standard. As PMM2 is a cytosolic protein and was depleted in our internal standard, it was excluded from further analysis. Furthermore, to enable monitoring of all remaining 29 target proteins within one LC-MRM measurement, we selected 132 out of the aforementioned 181 peptides (on average, 4 – 5 peptides per protein) based on optimal signal-to-noise ratios.

The setup of the MRM assay with the Skyline software in combination with an isotopically labeled protein standard allowed the transfer of the whole workflow including sample preparation to another laboratory. A set of samples, in detail described below, was processed and analyzed within one week in the group of Oliver Schilling at the Albert-Ludwigs-University in Freiburg, with basically the same results as obtained in Heidelberg by us (Figure 3b).

### Relative Abundances of Glycosyltransferases in HEK 293T, HeLa and Skin Fibroblast Cell Lines

In a SILAC based proteomics analysis, sample and control are grown in parallel in the normal „light“ and the „heavy“ SILAC medium, before being mixed and analyzed. This procedure can be facilitated by the production of a heavy labeled control in tissue cell culture in large amounts, which is then added as a spike-in SILAC standard in appropriate amounts to each sample grown in normal medium.^29^ We now asked the question whether an enriched membrane fraction of HEK 293T cells can be used as an internal standard for the analysis of glycosyltransferases even with different human cell lines like the commonly used HeLa cells or primary human fibroblasts. Therefore, we took whole cell lysates from HEK 293T and HeLa cell lines as well as from three primary wild type human skin fibroblast cell lines that were considered biological replicates. After the addition of the internal standard, proteolytic digest and LC-MRM analysis, 22 target proteins involved in *N-*glycosylation as well as *O-* and *C-* mannosylation (Figure 2) could be consistently quantified. For most proteins, only minor differences in relative protein amounts (≤ 2-fold) were observed across the investigated cell lines, which means that the HEK 293T-derived internal standard is appropriate also for these other cell lines. Notably, all three members of the functional building block ALG1/ALG2/ALG11 were significantly upregulated by approx. a factor of two in human fibroblasts compared to HEK 293T cells, which may indicate coordinated protein complex assembly. To investigate the ALG1/ALG2/ALG11 building block further, we next analyzed fibroblast cells originating from CDG type I patients carrying mutations in *ALG1*, *ALG2* or *ALG11* genes.

**Figure 2.**
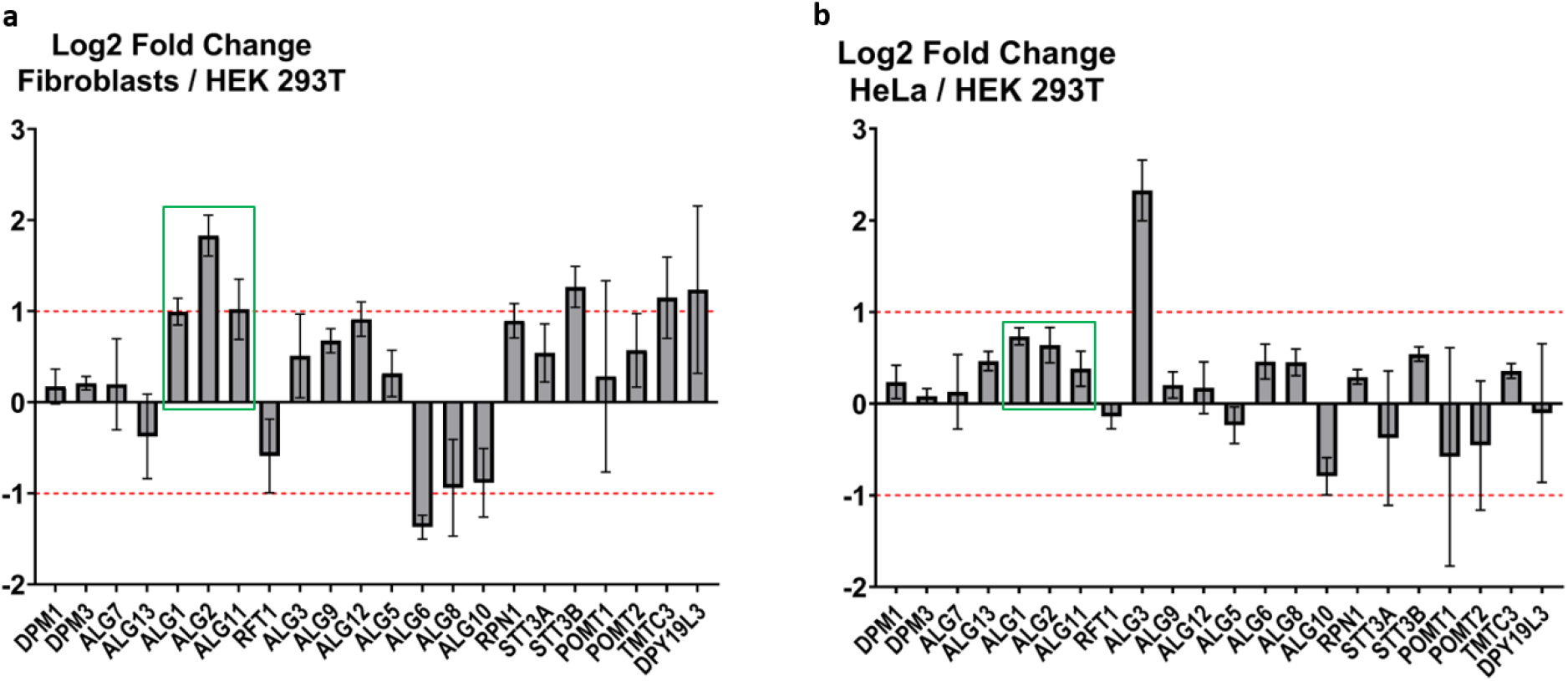
Relative quantification of a panel of 22 glycosyltransferases in human fibroblasts and HeLa cells compared with HEK 293T cells. Ratios between whole cell lysates and the internal standard were obtained and used to calculate fold changes between (A) fibroblasts and HEK 293T and (B) between HeLa and HEK 293T. Highlighted with a green box are changes in the functional unit ALG1, ALG2 and ALG11. Red dotted lines indicate the range between +/− two-fold change in relative protein abundance. Error bars show 95 % confidence intervals.

**Figure 3.**
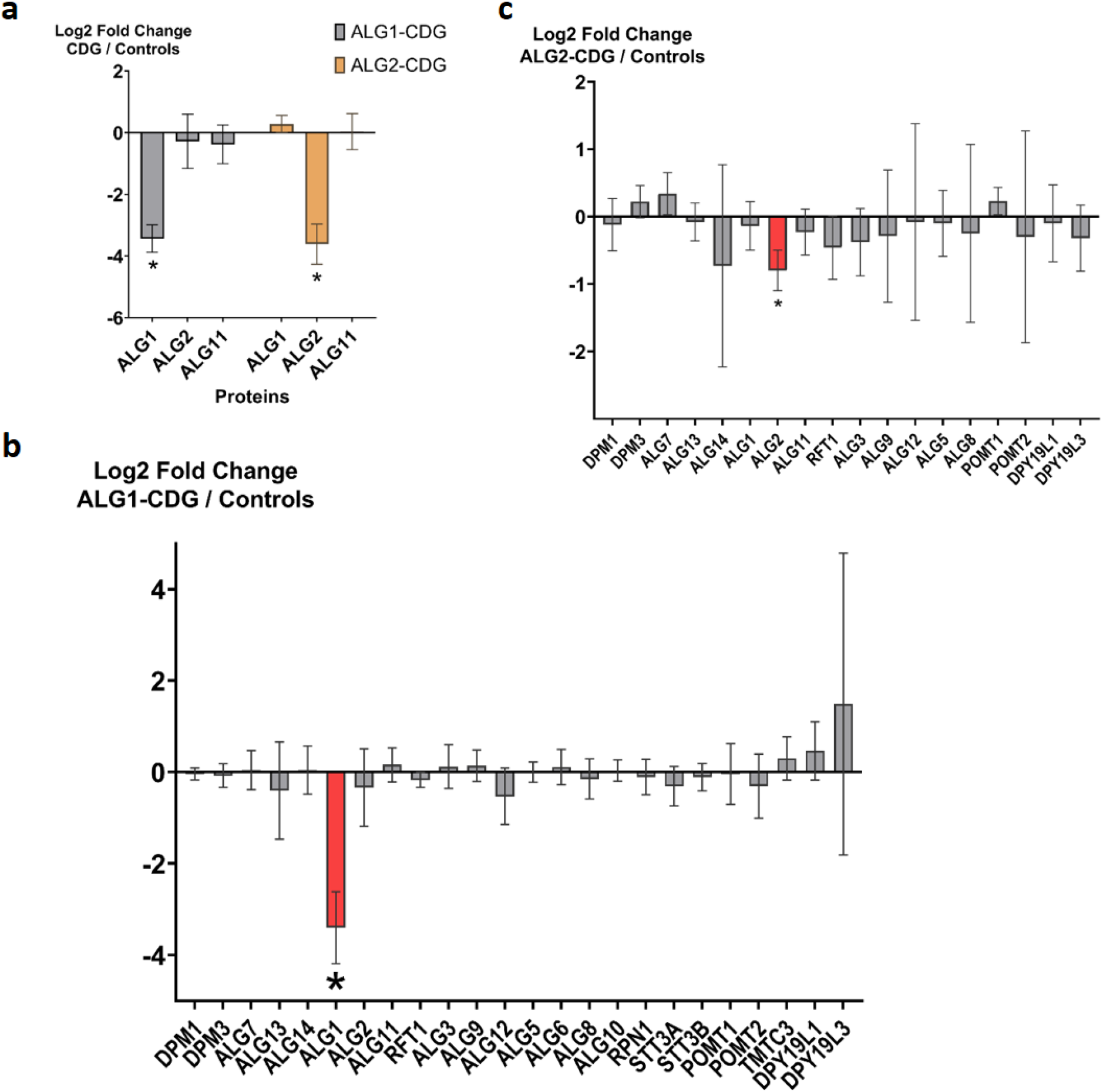
Comparison of ALG1/2/11 protein and transcript levels in fibroblast controls and ALG1/2-CDG patient samples. (a) ALG1 shows a strong decrease in ALG1-CDG patients without affecting ALG2 and ALG11. For the ALG2 patient ALG2 is decreased to similar extent without affecting ALG1 and ALG11. (b) LC-MRM results for ALG1-CDG patient fibroblasts compared to wild type controls acquired in Freiburg. Only ALG1 shows a significant and strong decrease, indicated with an asterisk and red color. (c) Comparison of mRNA transcript abundance between fibroblast controls and fibroblasts derived from an ALG2-CDG patient. Only mRNA levels of ALG2 are significantly decreased to approx. 60 % of wild type cells (adjusted p-value = 0.01). Error bars show 95 % confidence intervals.

### Comparison of Protein and mRNA Abundance between CDG Patient Fibroblasts and Controls

Since the membrane fraction of HEK 293T cells can be used as an internal standard for primary fibroblasts, we could ask the question about how levels of glycosyltransferases are affected in patients suffering from three subtypes of CDG with known mutations in *ALG1*, *ALG2* or *ALG11* genes. First, transcript levels from fibroblasts of altogether eight CDG patients^30–32^ were determined for a panel of 18 genes corresponding to the glycosyltransferases present in the MRM assay. No significant changes were observed compared to healthy controls (Figure S2) except for the ALG2-CDG patient that is described below in detail. Subsequently, the MRM assay was used to determine the protein levels of the glycosyltransferases. No significant changes were observed for any of the glycosyltransferases, except for those directly affected by the mutations. For the fibroblasts from all three ALG1-CDG patients and the ALG2-CDG patient, we observed a reduction of the corresponding protein to levels close to background (approx. to 1/8th of the wild type protein level) compared to four control primary fibroblast cell lines (Figure 3a). This was also shown by Western blot analysis (Figure S3). As mentioned above, these results were successfully reproduced in Freiburg for the ALG1-CDG subset of samples, demonstrating the robustness of our assay (Figure 3b and S4). The data presented demonstrate that the observed reduction in protein abundance was not due to transcript degradation and that there is no regulation at the transcriptional level to compensate for the loss of function.

As set out above, there was a significant change in transcript levels for the ALG2-CDG patient, but only for *ALG2* itself (Figure 3c). Its transcript abundance of *ALG2* decreased to approx. 60 % of the levels in wild type cells, which is in agreement with previously published data^32^ (Figure 3c). This is, however, a small change compared to the strong reduction on ALG2 protein level. The data presented demonstrate that the observed reduction in protein abundance was not due to transcript degradation and that there is no regulation at the transcriptional level to compensate for the loss of function.

In contrast, ALG11 was clearly detected in all ALG11-CDG patients. ALG11 levels were reduced to 50 % of wild type levels for primary fibroblasts from three CDG patients with different ALG11 alleles (referred to as ALG11_I-CDG) and remained unchanged in fibroblasts from the fourth patient analyzed (referred to as ALG11_II-CDG) (Figure 4a). For the latter patient, ALG11-CDG is caused by a heterozygous single nucleotide variant leading to an amino acid substitution within one of the ALG11 peptides monitored by our MRM assay.^31^ Ratios of three ALG11 peptides showed no difference compared to control fibroblasts whereas the fourth peptide carrying the site of amino acid exchange was reduced to approx. 35 %, which may represent the translation of transcripts from the unaffected allele (Figure 4b). Furthermore, an MRM chromatogram trace that was present in the respective ALG11 patient sample but not in controls was shown to coelute with a synthetic heavy labeled peptide carrying the amino acid substitution in question (Figure S5).

**Figure 4.**
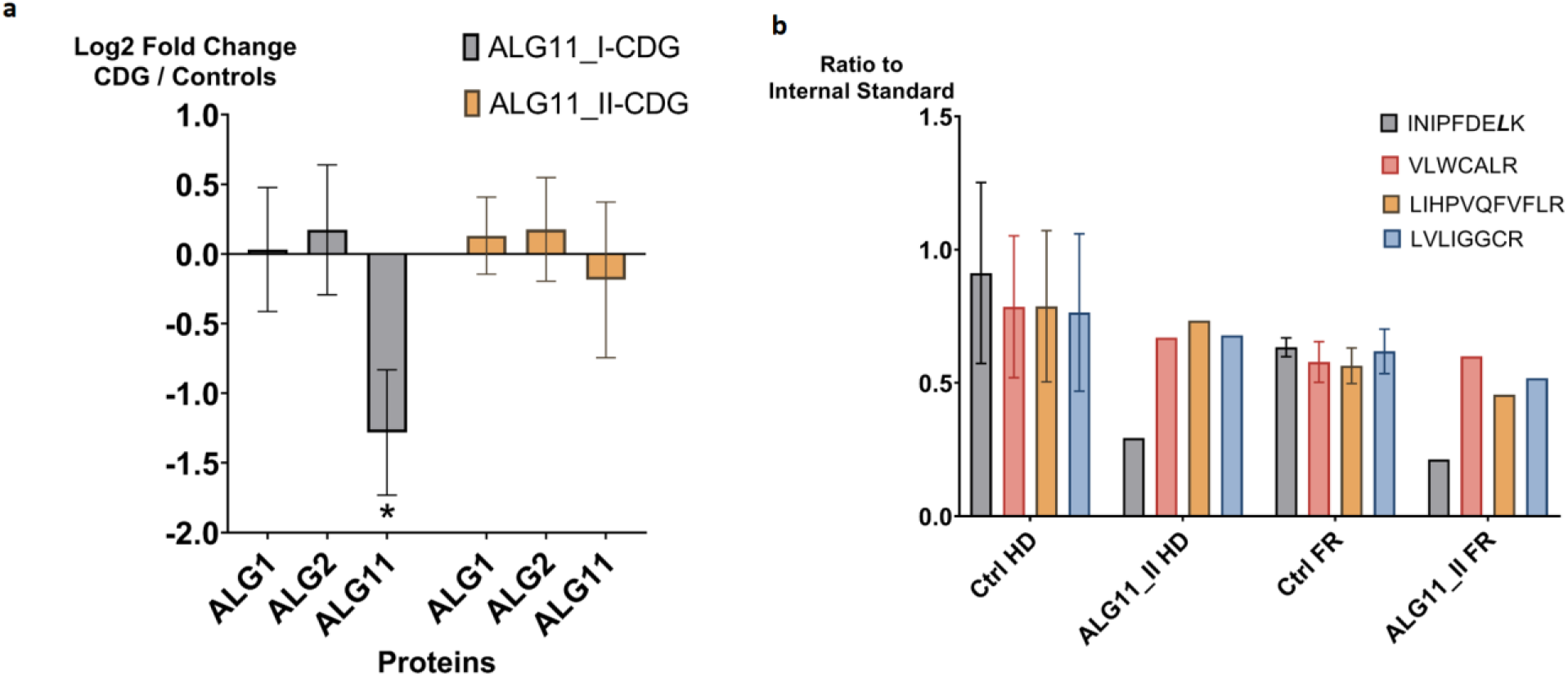
Comparison of ALG1/2/11 protein levels in fibroblast controls and ALG11-CDG patient samples. (a) In the ALG11_I-CDG group (three patients), ALG11 levels are significantly reduced to approx. 50 % of the control level. (b) For the ALG11_II-CDG patient, ALG11 protein is reproducibly quantified by 4 peptides in Heidelberg (HD) and Freiburg (FR). One of the peptides harbors an amino acid substitution site (L, italics) and shows a decrease in signal compared to the other three peptides.

ALG1, ALG2 and ALG11 form a functional unit and were shown to assemble into a complex in yeast.^6^ We expected that loss of one member of the protein complex also destabilizes other members as described for the OST complex in yeast.^33^ Strikingly, the almost complete loss of ALG1 in the CDG patients did not lead to a corresponding decrease of ALG2 and ALG11, and the same was observed for the ALG2 patient, when we looked at the ALG1 and ALG11 levels.

## Discussion

Most glycosyltransferases involved in glycosylation events in the ER are present at very low levels, which cannot be reliably quantified by discovery-based proteomics or antibody-based approaches. Even employing targeted proteomics approaches remains challenging when monitoring complete pathways comprised of low abundance membrane proteins.^34^ We established an MRM assay for 29 key glycosyltransferases, to get more information on the organization and regulation of the enzymes involved in the production of the Dolichol linked *N-*glycan precursor in the ER in humans. More than 20 of them are quantified across different experiments from crude cell lysates of HEK 293T and HeLa cells and skin fibroblasts. The MRM assay development was facilitated by commercially available and affordable crude labeled synthetic peptides that ensured signal specificity and enabled the calculation of laboratory-independent iRTs relative to the widely-used iRT peptide set.^20^ Raw data were processed in a semi-automated way using Skyline and R, including objective and transparent parameters for data preprocessing mostly based on rdotp and symmetry scores and an available R-script for batch statistical data analysis. The workflow we present here for the development of MRM assays should be broadly applicable to various sets of proteins. Furthermore, the efficiency of this workflow suggests that the development of MRM assays is no longer a limiting step for the use of targeted proteomics.

For precise quantification, we describe the production of an isotopically labeled protein standard, which is by far more suitable for endogenous signal intensity normalization compared to spike-in peptide standards.^35, 36^ This enriched membrane fraction from HEK 293T cells can also be used for other human cell lines like HeLa, HEK 293T and human fibroblast cell lines, since the glycosylation pathways in the ER are conserved also in terms of abundance of the different glycosyltransferases. Another advantage of the enriched and heavy labeled membrane fraction is that a heavy to light ratio close to one for the target proteins is achieved without increasing the sample complexity much. Calculated from total protein amount, only about 20 to 28 % in any given experiment came from the internal standard. Thus, we were able to inject higher amounts of our sample material, which is beneficial when dealing with low level targets of interest. Nevertheless, heavy labeled whole cell lysate from HEK 293T cells can also be used as an internal standard, although with reduced data quality.

The combination of iRTs and the internal protein standard that is introduced early during sample processing allows to share this method between laboratories and instrument platforms without great effort (Figures 4b). To demonstrate this, Patrick Bernhard from Oliver Schillings group in Freiburg reproduced our results from ALG1 and ALG11 patient samples. He performed the whole processing workflow starting from cell lysates while using his own equipment including LC and MS from different manufacturers compared with our laboratory. The overall time-investment, from receiving cell lysates until MRM data acquisition was completed, was less than a week, as the adaptation of the MRM method to the new setup was straightforward.

Quantification of the ALG proteins by shotgun proteomics results in high variability. In HeLa cells, the range between the lowest and the highest number often exceeded one order of magnitude and it was the same when comparing their abundances between HEK 293T and HeLa cells (Figure S6). The combination of MRM and the internal protein standard allowed the detection of differences of less than a factor of two with narrow confidence intervals. For data normalization SEC63, a member of the translocon complex, was chosen as an abundant ER marker membrane protein. For most of the glycosyltransferases, abundances in the ER were quite similar between different cell types like the hypotriploid HEK 293T, the tumor cell line HeLa and wild type fibroblasts despite their different functions. It is, however, important to note that ALG1, ALG2 and ALG11 are upregulated by a factor of two in fibroblasts compared to HEK 293T. These three enzymes form a functional unit attaching the first five mannose residues to the N-glycan precursor at the cytosolic side of the ER, and in yeast it was shown that they form heteromeric protein complexes. This coordinated upregulation in human fibroblasts is a hint that these three enzymes are also assembled into a protein complex in humans. ALG6, ALG8 and ALG10, which add the glucose residues at the luminal side, are similarly downregulated in a coordinated way.

We analyzed fibroblasts from CDG patients carrying mutations in both alleles of ALG1, ALG2 or ALG11. Surprisingly, only slight changes in transcript levels encoding these glycosyltransferases were detected, if at all. On protein level, only glycosyltransferases directly corresponding to the genetic defect were affected. For ALG1-CDG and ALG2-CDG fibroblasts, respectively, the corresponding protein levels were reduced close to the background level. There is obviously no fail-safe mechanism available to counteract the loss of a member within these early steps of protein glycosylation.

The four ALG11-CDG patients showed a different pattern. The ALG11 protein abundance was either unaffected or reduced to 50 % of wild type levels (for homozygous as well as heterozygous mutations). Reduction to 50 % can be interpreted by transcripts from only one allele leading to a stable protein product. This protein is affected in enzymatic activity in the CDG patients. The fact that a stable protein product is detected in significant amounts in all cases analyzed is a strong hint that structural integrity of ALG11 is even more important than its enzymatic function. Importantly, for the ALG11_II-CDG patient with unchanged ALG11 abundance, the original case report^31^ described severe clinical characteristics. In addition, there was an even more pronounced accumulation of shortened oligosaccharide intermediates, which serve as ALG11 substrates in wild type cells, compared to ALG11_I-CDG patients. These facts indicate that not protein stability but rather the enzymatic activity of ALG11 was affected by the described *ALG11* mutations, and that the severity of clinical symptoms does not correlate with ALG11 abundance. Furthermore, fibroblasts from the ALG11_II-CDG patient carry one heterozygous mutation leading to a single amino acid replacement of leucine to serine in one of our target peptides. Whereas the relative amount of 3 of the 4 monitored ALG11 peptides remained unchanged, the amount for the peptide that carried the leucine of the *wt* sequence was reduced to 35 % of wild type levels. The presence of the mutated peptide was proven in a separate experiment. Although a lucky coincidence, such a result could not have been found by immunoblotting. Importantly, this effect was also precisely reproduced in Freiburg and emphasizes the accuracy of our quantification.

*N-*glycosylation is highly conserved between yeast and humans^37^ and there is co-immunoprecipitation-based evidence that in yeast^6^ ALG1, ALG2 and ALG11 form a heteromeric complex, with ALG1 as the central subunit. If there is such a complex in humans, our data show that the loss of ALG1 does not destabilize the other components of the complex because ALG2 and ALG11 protein levels did not show any significant change, as one could expect from complex subunits and as it was demonstrated for the OST complex in yeast by MRM.^33^ The same is true for the loss of ALG2, as ALG1 and ALG11 remained unchanged. It is, however, different for ALG11. None of the mutations of four patients reduced protein levels to less than 50 % of the wild type protein, which may point in the direction that in the case of ALG11 loss of protein structure is even more severe than loss of activity and no viability may be possible.

Taken together, our MRM assay allows us to comprehensively study glycosylation in the ER by quantifying glycosyltransferases from complex biological samples. We are providing a new tool to support future research in this field and, possibly, to advance challenging diagnostic procedures for CDGs, for example when point mutations are linked to disease.

## Acknowledgments

We thank Ronny Heidasch, Julia Knopf and Marius Lemberg for providing HEK 293T cells and for excellent support for membrane preparation and Anne-Lore Schlaitz for kindly providing HeLa cells. We thank Oliver Schilling for the opportunity to perform MRM measurements at Freiburg University. We are grateful to Sabine Strahl, Sebastian Schuck and Jeroen Krijgsveld for help, critical discussions and feedback.

Funded by the Deutsche Forschungsgemeinschaft (DFG, German Research Foundation) –Project-ID FOR2509 P09 and P05, – Forschungsgruppe FOR 2509.

The authors declare no competing financial interest.

