## Supplementary Material for "Targeted Proteomics Reveals Quantitative Differences in Low Abundance Glycosyltransferases of Patients with Congenital Disorders of Glycosylation"

* equal contribution
### corresponding author

| | |
|---|---|
| Materials and Methods: | Page S-2 |
| Supplementary References: | Page S-4 |
| Supplementary Table 1 | Page S-5 |
| Supplementary Figures 1 – 6 | Page S-6 |

### Supplementary Materials and Methods

#### Western Blot Analysis

For Western blot 10 µg of total protein derived from patient and control fibroblasts were used. Samples were mixed with 6x Laemmli buffer (375 mM Tris–HCl, pH 6.8, 6 % SDS, 48 % glycerol, 9 % 2-mercaptoethanol, 0.03 % bromophenol blue) and denatured at 95 °C for 5 min. Extracts were analyzed on a 12.5 % SDS-PAGE and blotted onto a nitrocellulose membrane (GE Healthcare) by semi-dry electrophoretic transfer. The membrane was blocked for 1 h at room temperature (RT) with 5 % milk powder in PBST (0.1 % Tween20 (SERVA) in PBS). After blocking, the membrane was washed and incubated with the primary antibodies against ALG1 (Proteintech, 12872-1-AP, 1:1000 dilution) or ALG2 (Invitrogen, PA5-43263, 1:1000 dilution) overnight at 4 °C. After washing, the membrane was incubated with secondary antibody anti-rabbit IgG-conjugated with horseradish peroxidase (Santa Cruz; 1:10,000) for 45 min at RT. Protein signals were detected by light emission with Pierce™ enhanced chemiluminescence reagent (ECL) Western blot analysis substrate. After stripping the blots with 10 % acetic acid for 15 min the membrane was blocked again with 5 % milk powder in PBST for 1 h. Membranes were incubated with primary antibodies against ß-actin antibody (1:10,000, A5441, Sigma) for 1 h at room temperature. The membrane was subsequently incubated with secondary antibody anti-rabbit IgG-conjugated with horseradish peroxidase for 45 min and detected with a Pierce ECL plus assay kit (Thermo Fisher Scientific). All experiments have been repeated a minimum of three times. Western blot data was normalized to ß-actin signal and controls were set to 100 %.

#### Membrane Protein Fraction Preparation

The preparation of the membrane protein fraction was performed as previously described.[1] The hypotonic buffer contained 8 µM pepstatin, 20 µM leupeptin, 1 mM aprotinin and 1 mM phenylmethylsulfonyl fluoride as protease inhibitors; the post-nuclear supernatant was centrifuged at 100,000 g for 30 min at 4 °C through a low-salt sucrose cushion (500 mM sucrose, 50 mM HEPES-KOH, pH 7.4, 50 mM KOAc, 5 mM $Mg(OAc)_2$) and the last centrifugation was performed at 140,000 g for 30 min at 4 °C through an alkaline sucrose cushion as described.

#### Peptide Candidate Selection Procedure

Potential peptides resulting from an *in silico* digest in Skyline were selected due to the following criteria: tryptic, no missed cleavages, 7-21 amino acids in length and no methionine residues. Reported natural variants due to the UniProtKB database[2] (visualized with Protter[3] via a Skyline plugin) were avoided, if possible. Furthermore, selected peptides were checked for uniqueness for the respective protein against the human canonical background proteome, retrieved from the UniProtKB database (www.uniprot.org, Proteome ID: UP000005640, downloaded on 16.04.2018). In addition, the availability of already existing mass spectrometric (MS) data for each respective peptide was checked by using our own DDA data as well as the publicly available repository SRMAtlas (www.srmatlas.org)[4]. When little peptide candidates were available due to the amino acid sequence, methionine-containing peptides could be further considered if observed before in our own DDA data (at the end, 8 out of 181 peptides contained one or two methionines). Whenever still presented with more than ten remaining peptide candidates after aforementioned filtering steps, the predicted detectability of each peptide was checked by using the CONSeQuence web interface from the University of Manchester (http://king.smith.man.ac.uk/CONSeQuence),[5] where a score of 3-4 was considered as a detectable peptide. The merged information from the mentioned sources were used as basis for the selection of the most promising peptides, which can serve as proxy for the protein of interest in the MS context.

**MRM-Initiated Detection and Sequencing (MIDAS) Settings and Data Analysis**

MIDAS was used to identify appropriate MRM transitions for the heavy labeled synthetic peptides (1 pmol per injection) as described.[6] LC-MIDAS measurements were performed on the QTRAP® 5500 (Sciex) system in the positive ion mode with 50 maximum transitions per sample injection. Briefly, we used 30 ms dwell times for unscheduled MRM, a mass tolerance of 0.25 Da and an intensity threshold of 3000 counts/s, followed by up to two Enhanced Product Ion scans in the range of 230 – 1000 Da. Dynamic exclusion was set to 9 s. Due to the high abundance of the iRT peptides, 9 of the 10 iRT peptides were set to the permanent exclusion list, whereas the one remaining iRT peptide (DGLDAASYYAPVR) was used as positive control for the following database search. Resulting MS2 spectra were further preprocessed using the mascot.dll script within Analyst® 1.6.2 (Sciex) to generate an mgf-file. 2+ and 3+ were used as default precursor charge states, peaks with intensity of less than 1 % of the maximum intense peak were removed and spectra with less than 10 peaks were rejected. The generated mgf-file was then used to independently and objectively identify the respective precursors by performing a Mascot Database Search against the aforementioned canonical and isoforms containing protein database extended by common contaminants. The database search was performed with trypsin as the used digestion enzyme, with no allowed missed cleavages, a precursor (MS1) tolerance of 1 Da and a fragment (MS2) tolerance of 0.3 Da. Only doubly and triply positively charged precursor ions were allowed. Carbamidomethylation of cysteine was selected as a fixed modification. Methionine oxidation together with the selected labels for lysine 13C6 15N2 (+ 8 Da) and arginine 13C6 15N4 (+ 10 Da) were set as variable modifications. The results of the database search were exported as Mascot .dat files and inspected in Skyline as spectral libraries.

#### Supplementary Tables

**Table S1.** List of Used Cell Lines

| Cell Line | Origin | Source |
|---|---|---|
| HEK 293T | Human Embryonic Kidney | kindly provided by Dr. Marius Lemberg (ZMBH, Heidelberg) |
| HeLa CCL2™ | Human Cervical Cancer | ATCC, Manassas, USA; kindly provided by Dr. Anne-Lore Schlaitz (ZMBH, Heidelberg) |
| PCS-201-010 ™ (Control-A) | Human Skin Fibroblasts, neonatal | ATCC, Manassas, USA |
| SCC058 (Control-B) | Human Skin Fibroblasts | Merck, Darmstadt, Germany |
| CCD-39Sk (Control-C) | Human Skin Fibroblasts | ATCC, Manassas, USA |
| N21 (Control-D) | Human Skin Fibroblasts | Internal control cell line University Hospital Heidelberg; kindly provided by PD Dr. Christian Thiel (University Hospital Heidelberg) |

Table S2 – 4 are provided as sheets within a separate Excel file.

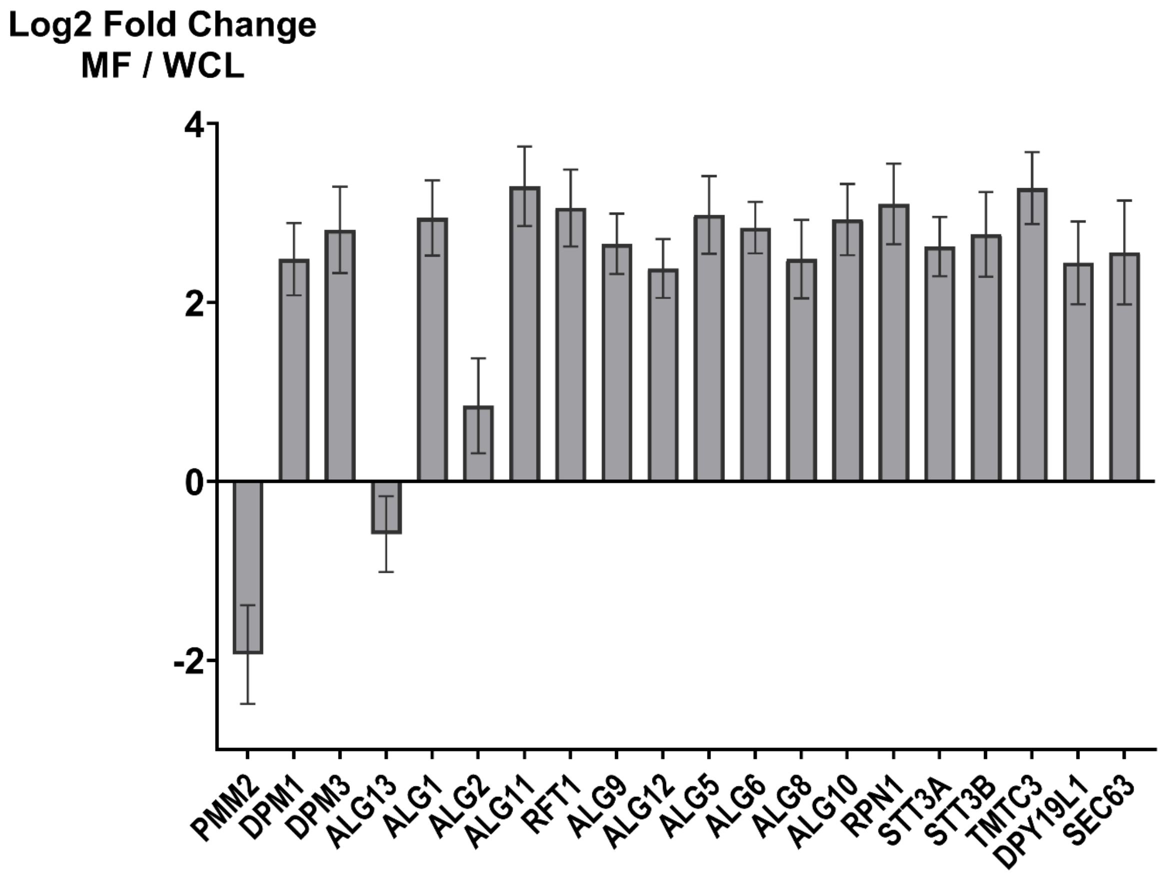


**Figure S1.** Comparative analysis of protein fold changes in HEK 293T whole cell lysate (WCL) and membrane fraction (MF). Isotopically labeled HEK 293T cells were either lysed with RIPA buffer to obtain WCL or by following the previously published protocol[1] to obtain an MF. Two independently performed MF preparations were compared to a WCL, in technical LC-MRM measurement triplicates, respectively (n=6 for MF and n=3 for WCL). 4 µg of total protein from the WCL or the MF were used for LC-MRM analysis in a randomized sample order. The resulting peak areas were normalized to 10 iRT peptides as global standards and protein fold changes were calculated in Skyline 20.1. Error bars show 95 % confidence intervals. All adjusted p-values were 0.01 for ALG13 or lower.

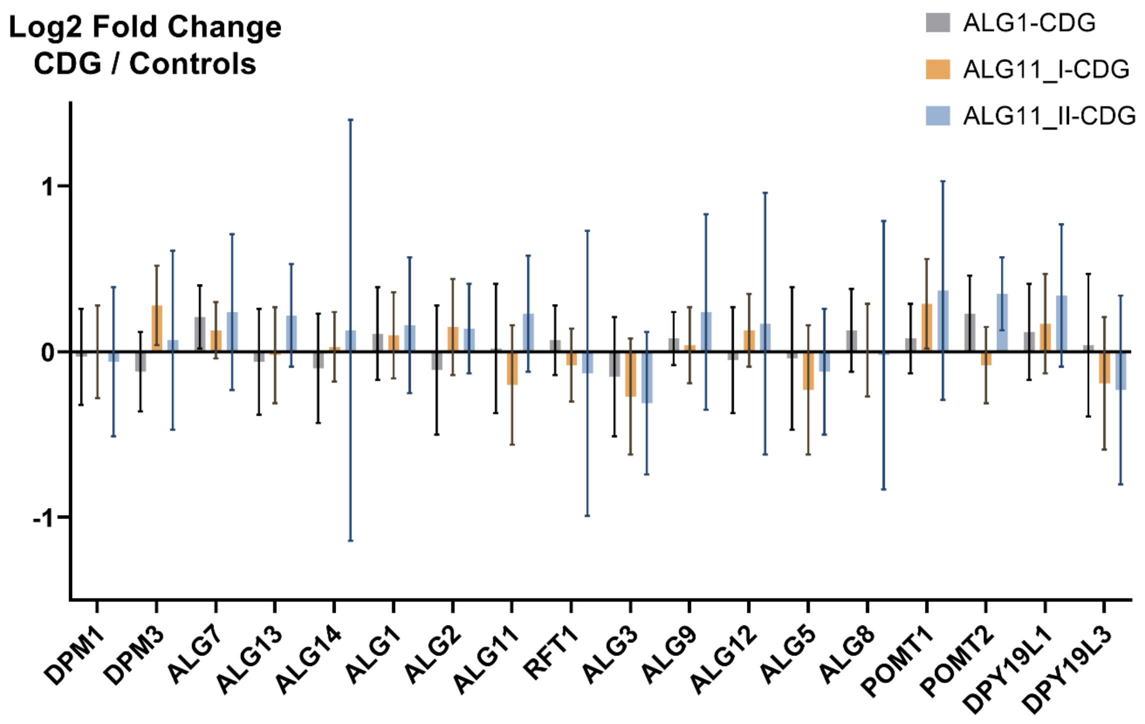


**Figure S2.** Comparison of mRNA transcript abundance between fibroblast controls and fibroblasts derived from CDG patients. No significant fold changes were detected. Error bars show 95 % confidence intervals.

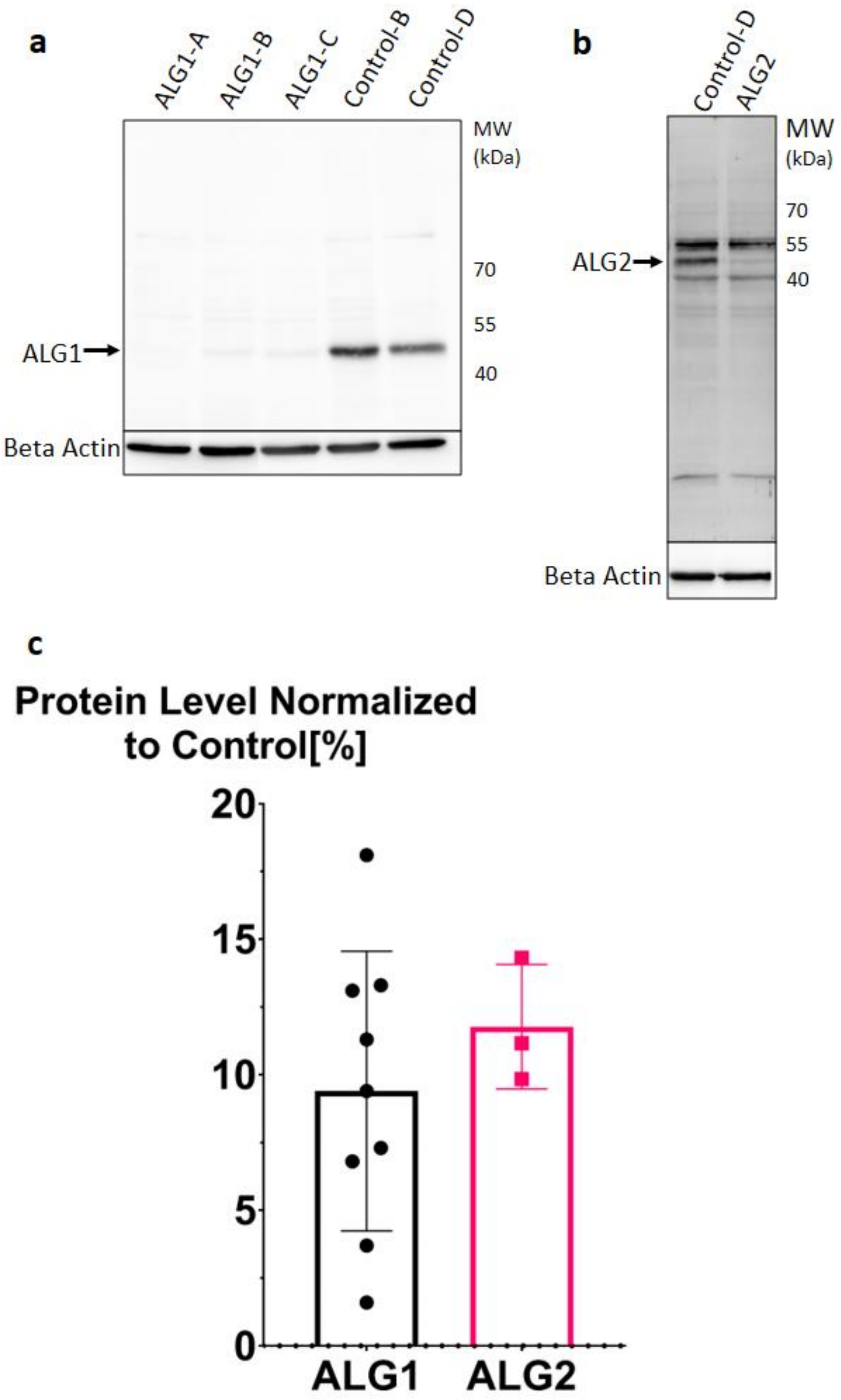


**Figure S3.** Validation of quantitative LC-MRM results by Western blot (WB) analysis. Representative blots for three ALG1-CDG patients and two control fibroblast cell lines (a) as well as for one ALG2-CDG patient and one control cell line (b) are shown. (c) Quantitative result summary for all WBs performed (n=3). Error bars show 95 % confidence intervals.

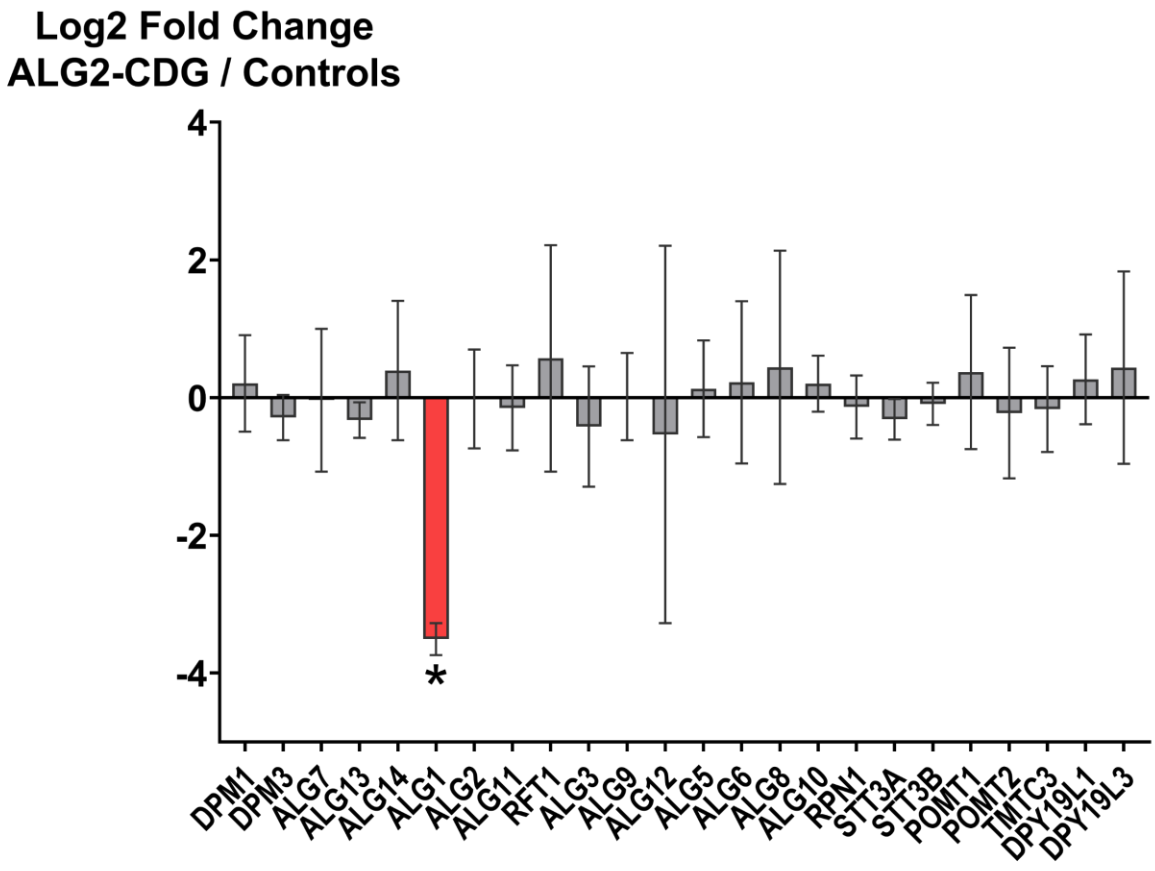


**Figure S4.** LC-MRM results for ALG1-CDG patient fibroblasts compared to wild type controls acquired in Heidelberg. Only ALG1 shows a significant (adjusted p-value < 0.0001) and strong decrease, indicated with an asterisk and red color. Error bars show 95 % confidence intervals.

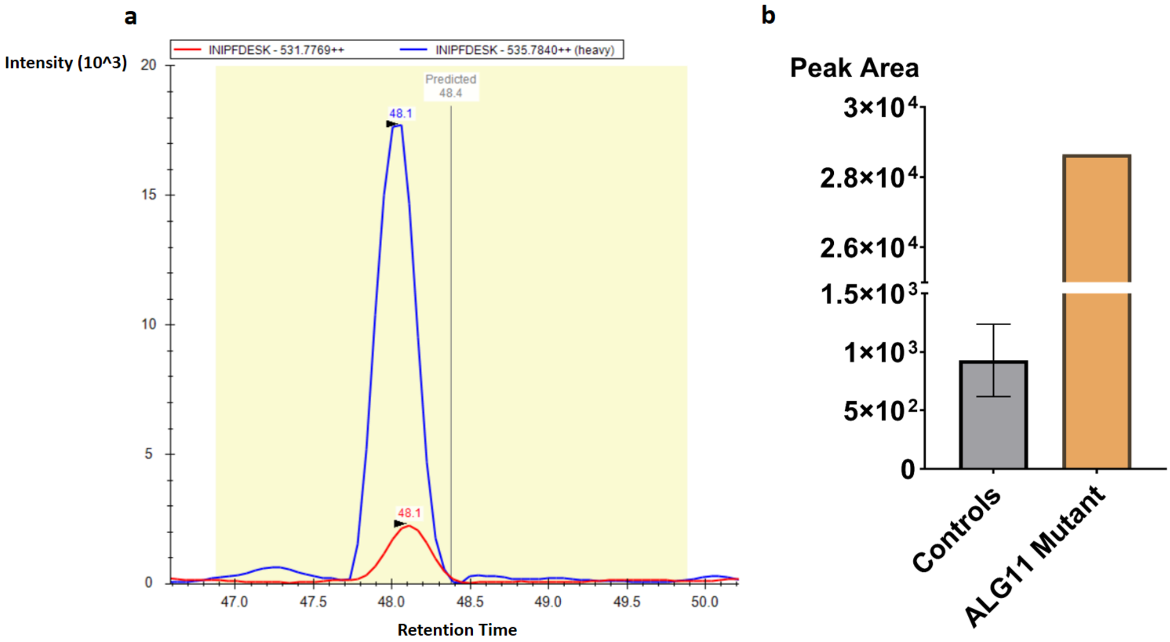


**Figure S5.** The peptide INIPFDESK containing an amino acid substitution is specifically detected in the respective patient ALG11-CDG-D. (a) Elution profile of the synthetic, heavy labeled peptide INIPFDESK (blue) validates the signal (red) found in the patient fibroblast sample in Freiburg as the mutated target peptide. The retention time was also excellently predicted by the iRT calculated in Heidelberg. (b) The validated signal from figure part (a) is present in the patient and essentially absent in controls. Error bars show standard error of mean.

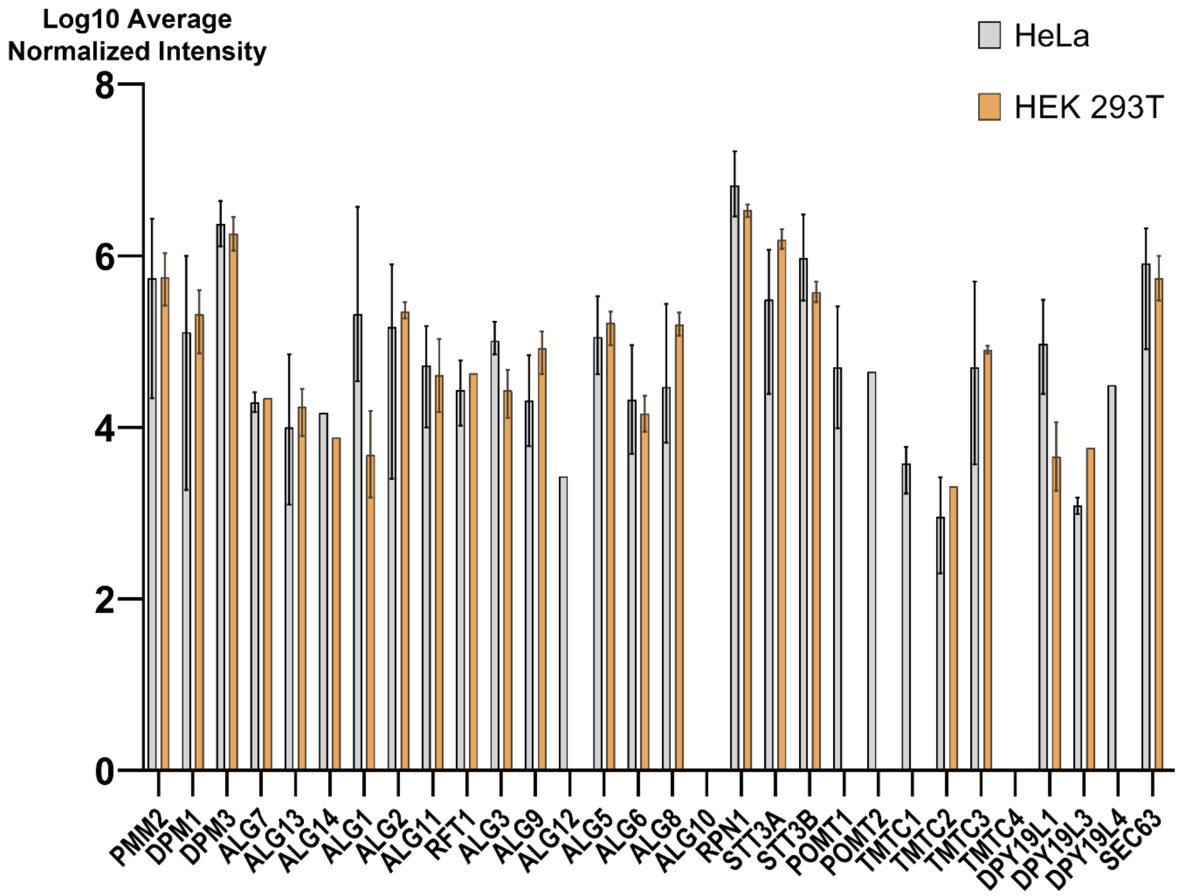


**Figure S6.** iBAQ values from the ProteomicsDB. Average values of glycosyltransferases are shown for HEK 293T cells and HeLa cells. Minimal and maximal values from Proteomics DB are indicated by error bars.
